# scProtoTransformer: Scalable Reference Mapping Across Molecules, Cells and Donors

**DOI:** 10.64898/2025.12.06.692735

**Authors:** Zhenchao Tang, Haohuai He, Shouzhi Chen, Jun Zhu, Tianxu Lv, Jiale Zhou, Jiehui Huang, Guanxing Chen, Linlin You, Calvin Yu-Chian Chen

## Abstract

The rapid accumulation of single-cell data has made it possible to comprehensively characterize biological systems at molecular, cellular, and donor levels. However, scalable reference mapping across different resolutions remains a major challenge in current research. Here, we propose **scProtoTransformer**, a prototype-based Transformer architecture designed to achieve scalable reference mapping across molecular, cell, and donor levels. scProtoTransformer introduces a knowledge-guided prototype tokenizer that projects gene expression into biologically interpretable pathway prototypes, effectively reducing numerical batch effects while preserving biological semantic patterns. Furthermore, by leveraging knowledge distilled from the foundation model and a dynamic supervised fine-tuning strategy, scProtoTransformer achieves robust biological representations with reduced pre-training requirements. Benchmark experiments across molecular, cell, and donor-level reference mapping demonstrate that scProtoTransformer delivers competitive or even superior performance compared with state-of-the-art approaches, while providing interpretability through biologically prototypes. Together, these results establish scProtoTransformer as a unified framework for scalable reference mapping, laying the foundation for systematic understanding from genes to individuals.

## 1 Introduction

The rapid development of single-cell measurement and analysis technologies is driving a comprehensive understanding of biological systems at the gene, cell, and tissue levels. The rapid accumulation of data has already made deep learning the most mainstream solution in the field [1]. Mapping annotations from reference datasets to query datasets remains a critical workflow in AI for Biology systems [2]. Moreover, reference mapping based on atlases can further enable numerous downstream applications [3–6]. Although many methods have been developed for reference mapping in cell identity annotation, scalable reference mapping across molecules, cells, and tissues remains an unexplored area. At the same time, multi-level representations are essential resources for building artificial intelligence–powered virtual cells [7].

However, achieving robust scalable reference mapping still faces two major challenges. The first is the classic problem: gene expression, as a numerical measurement, is highly susceptible to batch effects [8–10]. Differences in experimental platforms and sequencing depth can cause even the same cell type to exhibit significant discrepancies in expression profiles across datasets. Such “numerical drift” often obscures the true biological signals, leading models to learn technical noise rather than biological heterogeneity, thereby reducing generalization across datasets, platforms, and even laboratories. Thus, how to learn biologically meaningful cellular representations while avoiding interference from batch effects has long been a central challenge in single-cell analysis [11]. Many current integration methods rely on predefined batch information for computation, but in real-world scenarios, the sources of batch effects are often unclear, and subjective batch definitions may severely compromise the robustness of research conclusions [12].

The second challenge lies in the current focus of existing methods, which is limited to cell-level reference mapping. The most popular approaches employ pretrained foundation models to obtain cell representations and then use supervised fine-tuning to transfer labels from reference data to query data [13–17]. However, reference mapping across molecules, cells, and tissues has not yet been explored. Biological systems are inherently hierarchical [18]: at the molecular level, gene–gene interactions influence biological functions such as pathways; at the cellular level, different gene set patterns describe the heterogeneity among individual cells; at the donor level, phenotypes constructed by groups of cells capture inter-individual differences. To date, there is still no multi-level framework that can achieve representation and reference mapping simultaneously at the molecular, cellular, and donor levels [19]. We envision that such a framework would not only capture gene context at the molecular level, support cross-dataset integration and annotation at the cellular level, but also generalize cell group characteristics at the donor level, thereby enabling a systematic understanding from molecules to individuals.

To address these two challenges, we propose scProtoTransformer, a prototype-based Transformer architecture for scalable reference mapping across molecular, cellular, and donor levels. scProtoTransformer does not require subjective batch selection; instead, it abstracts gene expression into a set of biological prototypes, thereby mitigating the impact of numerical batch effects while retaining biological semantics. Specifically, inspired by prototype-based modeling in computer vision [20–22], we introduce the Prototype Tokenizer, which linearly projects gene expression vectors into a prototype space. The projection matrix of the Prototype Tokenizer is initialized with biological priors, ensuring that each prototype corresponds to a biologically well-defined pathway. These prototypes are then treated as tokens and fed into stacked Transformer layers, where the self-attention mechanism models interactions among prototypes, capturing cross-functional biological associations while avoiding interference from raw expression values.

In addition, we design a knowledge distillation strategy guided by foundation models for scProtoTransformer. The goal is to enable scProtoTransformer to inherit the knowledge obtained by foundation models at great training cost, without requiring large-scale pretraining itself. Specifically, we employ a frozen foundation model (e.g., scGPT) as a navigator, and use its cell embeddings to supervise scProtoTransformer’s learning. Unlike the unsupervised pretraining strategy used by foundation models, our approach can dynamically leverage available labels: when labels are present, their discriminative power enhances scProtoTransformer’s biological representation capacity. To further train the model on limited data, we use Dynamic SFT, which adaptively controls gradients to prevent overfitting.

Benchmark experiments demonstrate that scProtoTransformer supports multi-level reference mapping: the Prototype Tokenizer enables molecular-level reference mapping through gene embeddings; cell embeddings can be used for cell-level reference mapping; and aggregating embeddings of all cells from the same donor sample into existing multi-instance learning modules allows for donor-level reference mapping. In summary, scProtoTransformer lays the foundation for integrative analysis across molecular, cellular, and donor levels.

## 2 Results

### 2.1 scProtoTransformer overview

The workflow of scProtoTransformer is illustrated in Figure 1a. For each input single cell, its gene expression profile is converted by the prototype tokenizer into multiple prototype tokens. Stacked Transformer layers then learn the interactions among these prototype tokens, modeling the associations between biological functions. The cls token from the final Transformer layer is used as the cell embedding. In addition, we distill scProtoTransformer from the foundation model scGPT, requiring pretraining on only a small subset of scBaseCamp [23] (2 million cells sampled from 100 million, reducing the pretraining scale to 2% of the full dataset). Unlike unsupervised pretraining of foundation models, when label information is available in the data, we enhance the biological representation capacity of scProtoTransformer through Dynamic SFT.

**Figure 1:**
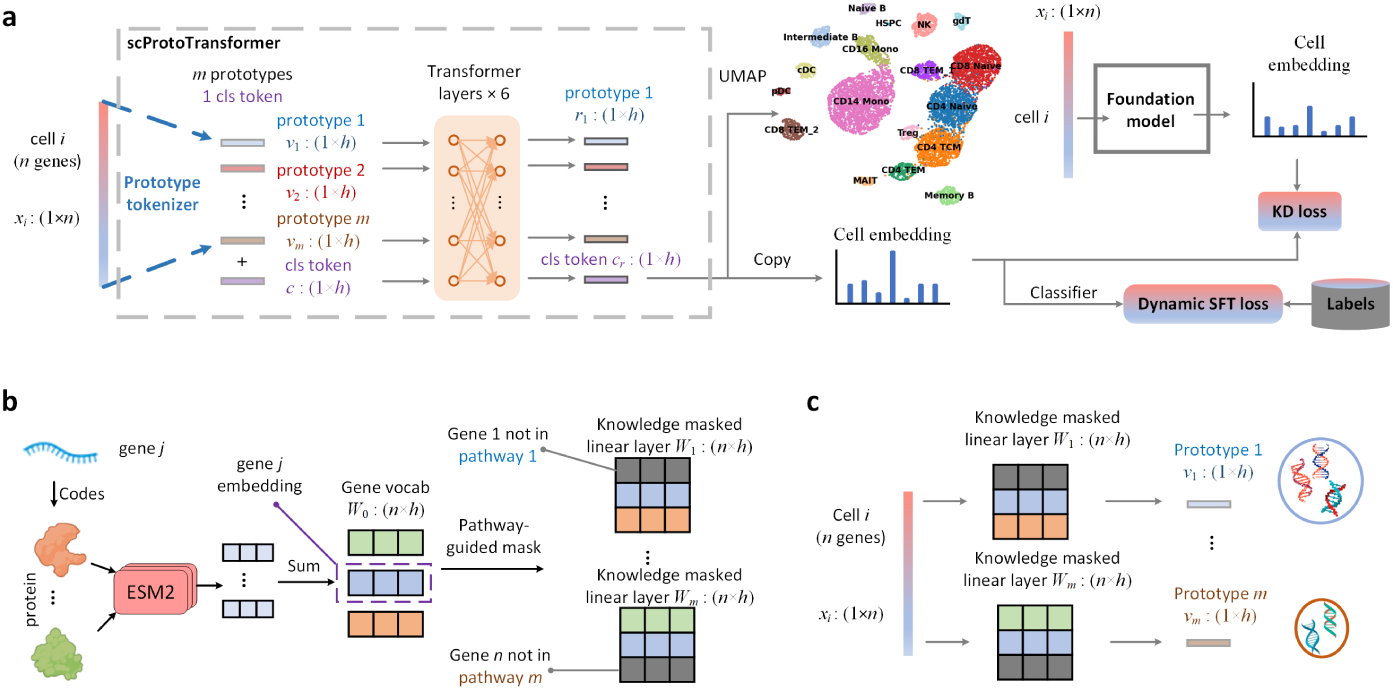
Architecture of scProtoTransformer. **a**: For a given single cell, the gene space is transformed into prototypes that represent biological functions. The self-attention mechanism learns the interactions among prototypes to model the associations between biological functions. During pretraining, we distill scProtoTransformer from a frozen foundation model. Instead of cross-entropy, we use a dynamic SFT loss that adaptively controls gradients to prevent model overfitting. **b**: Parameter initialization process of the prototype tokenizer. **c**: The prototype tokenizer converts the gene expression profile of a single cell into multiple parallel prototype representations.

More specifically, scProtoTransformer incorporates knowledge-based parameter initialization in the prototype tokenizer. As shown in Figure 1b, we first construct a gene vocab, where each row represents a gene embedding. Each gene embedding is obtained by summing the embeddings of proteins encoded by the corresponding gene. Next, we apply a pathway-guided mask to transform the gene vocab into m parallel linear layers. As shown in Figure 1c, these parallel linear layers convert gene expression into m prototype tokens.

After pretraining, each row in the gene vocab, i.e., the gene embeddings (stored using torch.nn.Embedding), supports molecular-level reference mapping. The cell embeddings produced by scProtoTransformer are used for cell-level reference mapping. For donor-level reference mapping, all cell embeddings from the same donor sample are aggregated through a multi-instance learning module to obtain the donor embedding.

### 2.2 Molecular level reference mapping

We extracted the gene vocabulary from scProtoTransformer and visualized the embeddings of 30,000 human genes using UMAP [24]. For comparison, we also extracted the gene embeddings from the foundation model scGPT. Using the labels provided in GenePT, we annotated each gene with its functional category. As shown in Figure 2a, the UMAP visualization colored by gene function types reveals distinct clusters, indicating that scProtoTransformer encodes gene functions more clearly than the foundation model, while ensuring that the gene embeddings preserve key biological information.

**Figure 2:**
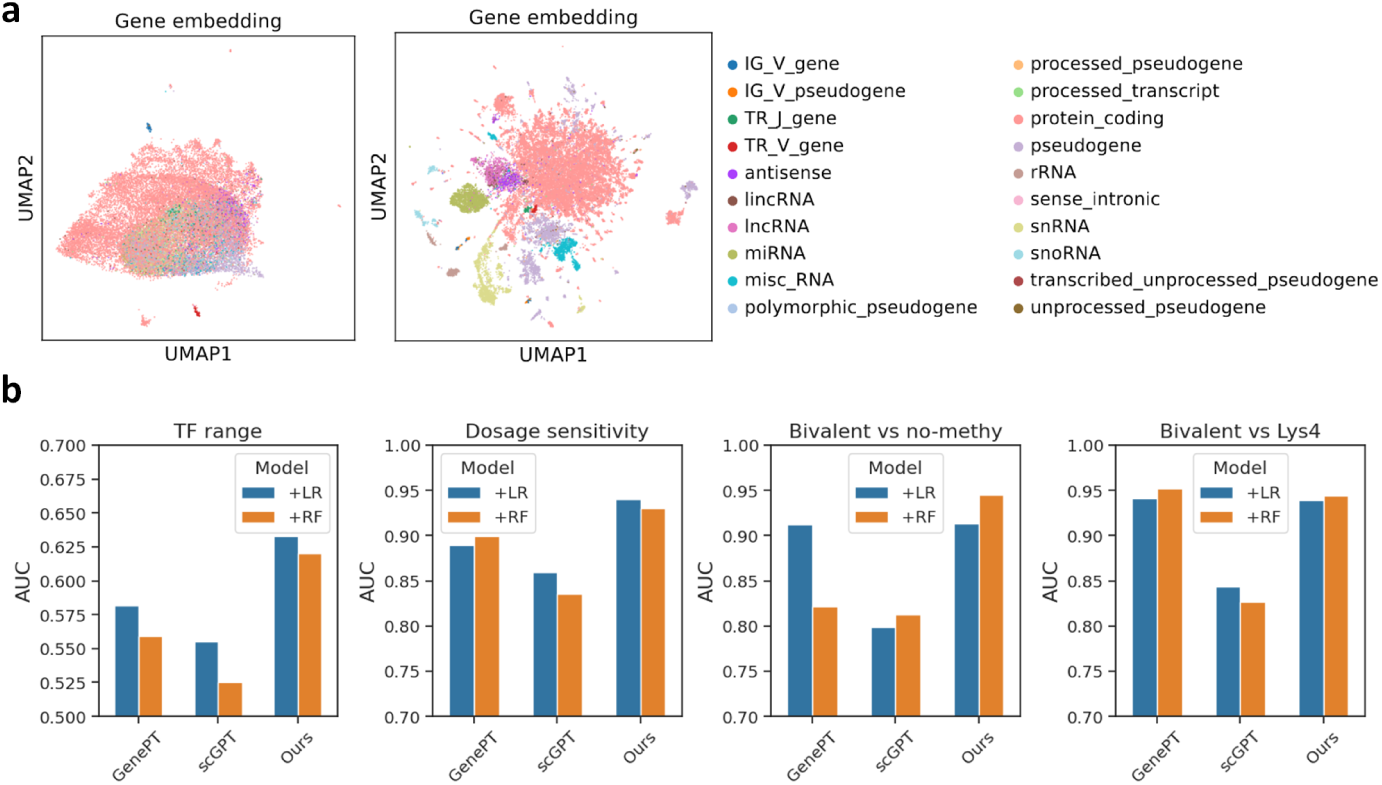
Molecular-level reference mapping. **a**: UMAP visualization of the human whole genome. The left shows gene embeddings from the foundation model, while the right shows our gene embeddings, colored by gene functional categories. **b**: Comparison of different methods across four molecular-level reference mapping tasks.

For gene-level reference mapping, we evaluated performance on specific biological tasks related to gene network dynamics. These tasks were drawn from datasets curated in the Geneformer benchmark [13]: long-range versus short-range transcription factors (TFs), dosage-sensitive versus dosage-insensitive TFs, bivalent versus non-methylated genes, Lys4-only-methylated versus non-methylated genes. For all four datasets, reference and query sets were partitioned according to the Geneformer protocol.

We compared our method with two mainstream approaches: GenePT [25] and scGPT [16]. For each method, we performed reference mapping using logistic regression (LR) and random forest (RF) classifiers trained on the corresponding gene embeddings. Performance was evaluated using the area under the receiver operating characteristic curve (AUC). As shown in Figure 2b, our method consistently achieves competitive results, despite GenePT and scGPT benefiting from large language models and large-scale pretraining, respectively. These results highlight the potential of scProtoTransformer for molecular-level representation.

### 2.3 Cell level reference mapping

Given a single-cell input, scProtoTransformer can output a cell embedding. We evaluated cell-level reference mapping on three cell type annotation datasets. Each dataset has distinct characteristics: The BMMC dataset [26] is measured with scRNA-seq and contains 13 batches and 22 cell types. The PBMC dataset [27] is measured with scATAC-seq and contains 2 batches and 6 cell types. Following Muse-GNN [28], we used Seurat to convert scATAC-seq peaks into scRNA-seq genes. The Pan-cancer dataset [29, 30] is measured with scRNA-seq but involves heterogeneous cell types across multiple cancers, making annotation more challenging. Pan-cancer includes 13 cancer types and 23 cell types, with each cancer treated as a separate batch.

Data preprocessing followed the standard pipeline of scanpy. For BMMC, we selected the first 6 batches as the reference dataset and the remaining 7 batches as the query dataset. For PBMC, we used the first batch as the reference dataset and the remaining batch as the query dataset. Pan-cancer was used to validate scProtoTransformer’s performance in cross-disease cell type annotation. Specifically, three cancers (ESCA: esophageal carcinoma, THCA: thyroid carcinoma, and UCEC: uterine corpus endometrial carcinoma) were used as the reference dataset, while the other cancers (batches) served as the query dataset.

We compared our method against the following annotation baselines: Seurat [31], CellTypist [32], ACTINN [33], TOSICA [34], Cellcano [35], MetaTiME [36], Geneformer [13], scBERT [14], CellLM [37], LangCell [38], scGPT [16], and KIDA [39]. For cell type annotation, we used accuracy (Acc) and F1 score as evaluation metrics. As shown in Figure 3, our method achieved superior cell type annotation results. Although foundation model-based methods generally performed well, their maintenance and data management are highly complex. Small models such as KIDA and TOSICA also showed strong performance, but they require retraining from scratch on each dataset, which poses a challenge for rapid deployment (training from scratch requires more complex hyperparameter design). By contrast, our method requires neither maintaining a complex foundation model corpus nor retraining from scratch, while still achieving competitive performance on cell-level reference mapping tasks.

**Figure 3:**
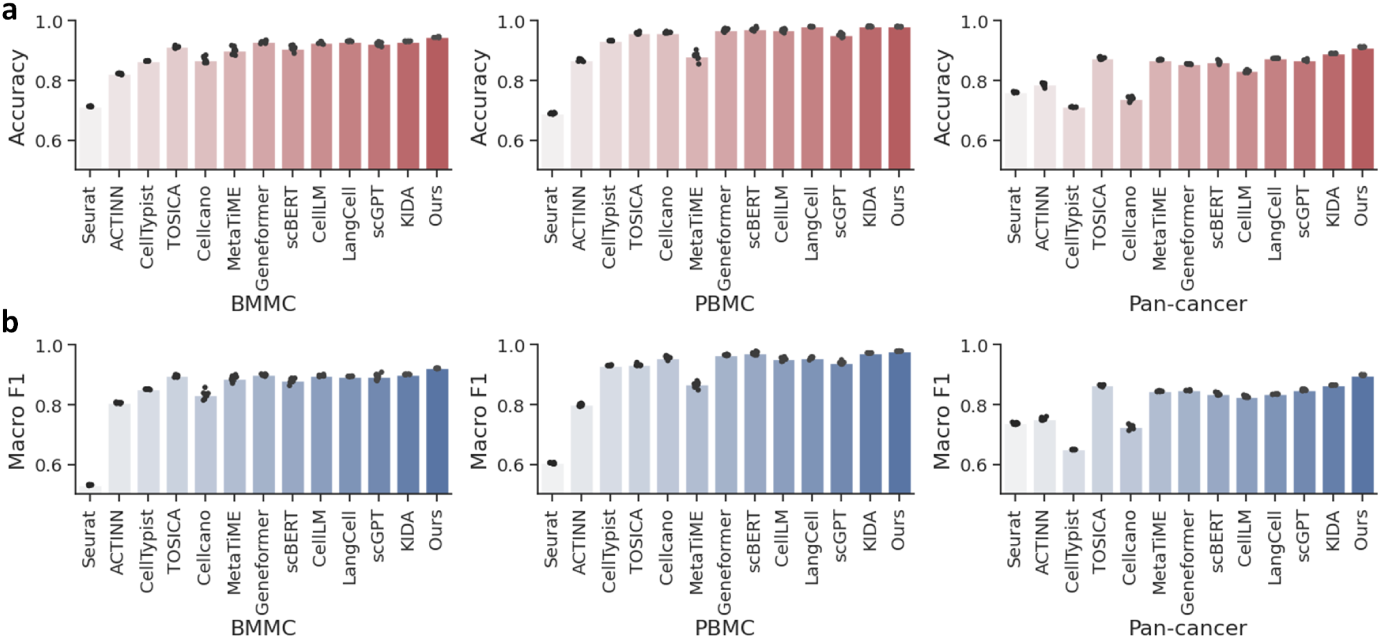
Cell-level reference mapping. **a**: Comparison of accuracy across different methods on cell-level reference mapping tasks. **b**: Comparison of F1 scores across different methods on cell-level reference mapping tasks.

For the two different modalities (BMMC: scRNA-seq, PBMC: scATAC-seq), we visualized the cell embeddings using UMAP. As shown in Supplementary Figure 1, compared with modality-specific methods and the original PCA, the cell embeddings generated by scProtoTransformer not only eliminated batch effects but also enhanced biological heterogeneity (the discriminability between cell types).

### 2.4 Explainability of prototypes

Cell-level embeddings exhibit strong biological heterogeneity, which can be analyzed through prototypes. scProtoTransformer provides inherent interpretability because each prototype can be named by a specific pathway. The input to each Transformer layer consists of a cls token, representing the cell embedding, along with a set of prototype tokens. Thus, each cell embedding can extract attention values associated with all pathways. By aggregating across all cells, we obtain an attention expression matrix. We then perform differential analysis on this matrix—for example, selecting the top 6 CD8+ T–specific pathways and visualizing them with a heatmap, as shown in Supplementary Figure 2a. Taking the top-ranked pathway as an example, we retained the genes involved in this pathway and examined their expression distribution across different cell types. We found that these genes are indeed highly expressed in CD8+ T cells (Supplementary Figure 2b), indicating that the attention learned by scProtoTransformer is consistent with the observed biological distribution, thereby demonstrating the interpretability of its prototype-based design.

Building on our previous strategy in KIDA for selecting top-ranked genes, we retained cell type–specific genes and constructed a gene co-expression network based on their embeddings. As shown in Supplementary Figure 3a, we first obtained 20 CD8+ T cell–specific gene embeddings. By calculating similarities among these embeddings, we generated a graph representing the CD8+ T–specific co-expression network. We identified five genes with particularly strong similarity. Visualizing their expression patterns with UMAP (Supplementary Figure 3b) confirmed that these genes are highly expressed in the CD8+ T cell region. Notably, when compared with CellMarker [40], all of these genes are known marker genes for CD8+ T cells.

### 2.5 Cross-modal and cross-batch integration

Since most downstream tasks benefit from a unified cell-level atlas, we further validated the representation capability of scProtoTransformer through integration tasks. We first evaluated multi-modal integration. We adopted the benchmark we previously established in Monae [41] and selected three challenging unpaired multi-modal datasets: 10xMultiome [42]: jointly profiled 9,631 human peripheral blood mononuclear cells (PBMCs) using 10X Multiome, with the pairing information between RNA and ATAC modalities removed. Ma2020 [43]: profiled 32,231 mouse skin cells with SHARE-seq to obtain RNA+ATAC modalities, with pairing information removed. Muto2021 [44]: profiled a total of 44,190 cells from the human kidney using snRNA-seq and snATAC-seq. The RNA modality contained 19,985 cells, and the ATAC modality contained 24,205 cells.

We compared scProtoTransformer against several specialized integration methods: GLUE [42], MultiVI [45], Monae [41], and Modal-NexT [46]. We used three widely adopted evaluation metrics in multi-modal integration: batch removal, biology conservation, and overall score. As shown in Figure 4a, scProtoTransformer achieved the best balance between batch effect removal and biological conservation. As shown in Supplementary Figure 4, scProtoTransformer outperformed state-of-the-art integration-specific methods in terms of overall score.

**Figure 4:**
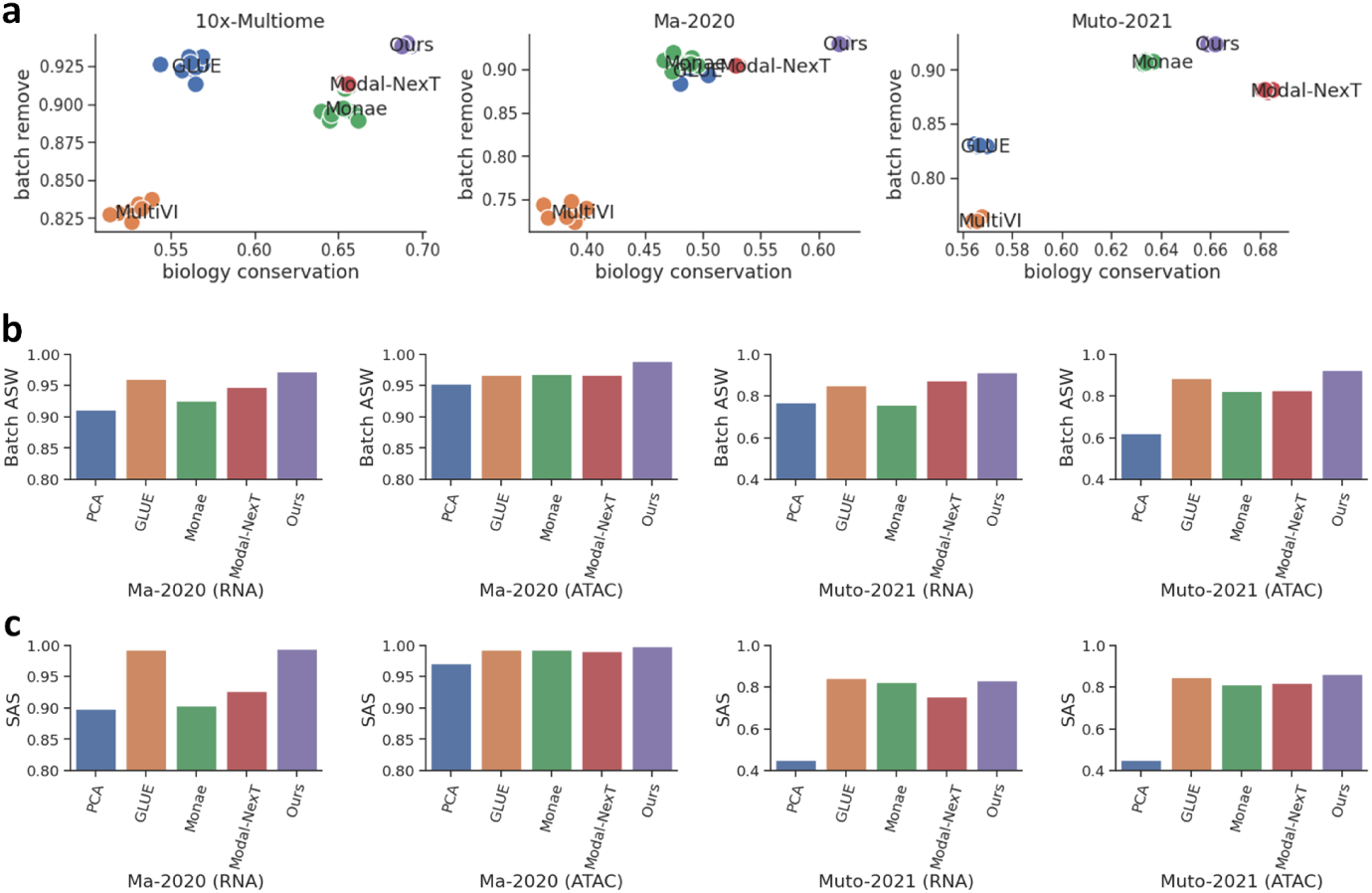
Comparison of multi-modal integration results and batch integration results. **a**: Simultaneous comparison of batch effect removal and biological heterogeneity preservation. **b**: Comparison of Batch ASW (Average Silhouette Width) across different batch integration methods. **c**: Comparison of SAS (Seurat Alignment Score) across different batch integration methods. In batch integration, PCA refers to cell embeddings without batch correction.

For Ma2020 and Muto2021, each modality consisted of 4–5 batches, enabling evaluation of batch integration metrics within each modality. The batch integration metrics include Batch ASW (Average Silhouette Width) and SAS (Seurat Alignment Score). As shown in Figures 4b–c, scProtoTransformer was able to automatically achieve batch integration during multi-modal integration, without requiring adversarial training strategies as in GLUE.

### 2.6 scProtoTransformer is suitable for spatial data

In multicellular organisms, cells within a tissue slice interact with each other and form spatial structural patterns, while batch effects are often more pronounced across different slices. Existing specialized spatial analysis methods typically require incorporating additional non-molecular features (e.g., spatial coordinates) to learn a joint representation across slices, highlighting the limitations of scRNA-seq analysis methods in handling spatial data. In contrast, scProtoTransformer has an inherent ability for batch integration, allowing it to be directly adapted to spatial data without relying on non-molecular features such as spatial coordinates, which can introduce subjective biases. We validated the adaptability of scProtoTransformer on the DLPFC dataset, which contains normalized mRNA expression and 2D spatial coordinates for each cell.

As shown in Figure 5a, UMAP visualization of PCA embeddings across multiple slices revealed strong batch effects, with cell embeddings from different slices completely separated. By contrast, UMAP visualization of scProtoTransformer cell embeddings (Figure 5b) showed complete mixing across slices, while clustering according to cell types could still be clearly observed. We compared scProtoTransformer with specialized spatial analysis methods using ARI as the evaluation metric, and as shown in Figure 5c, our method achieved superior representation quality.

**Figure 5:**
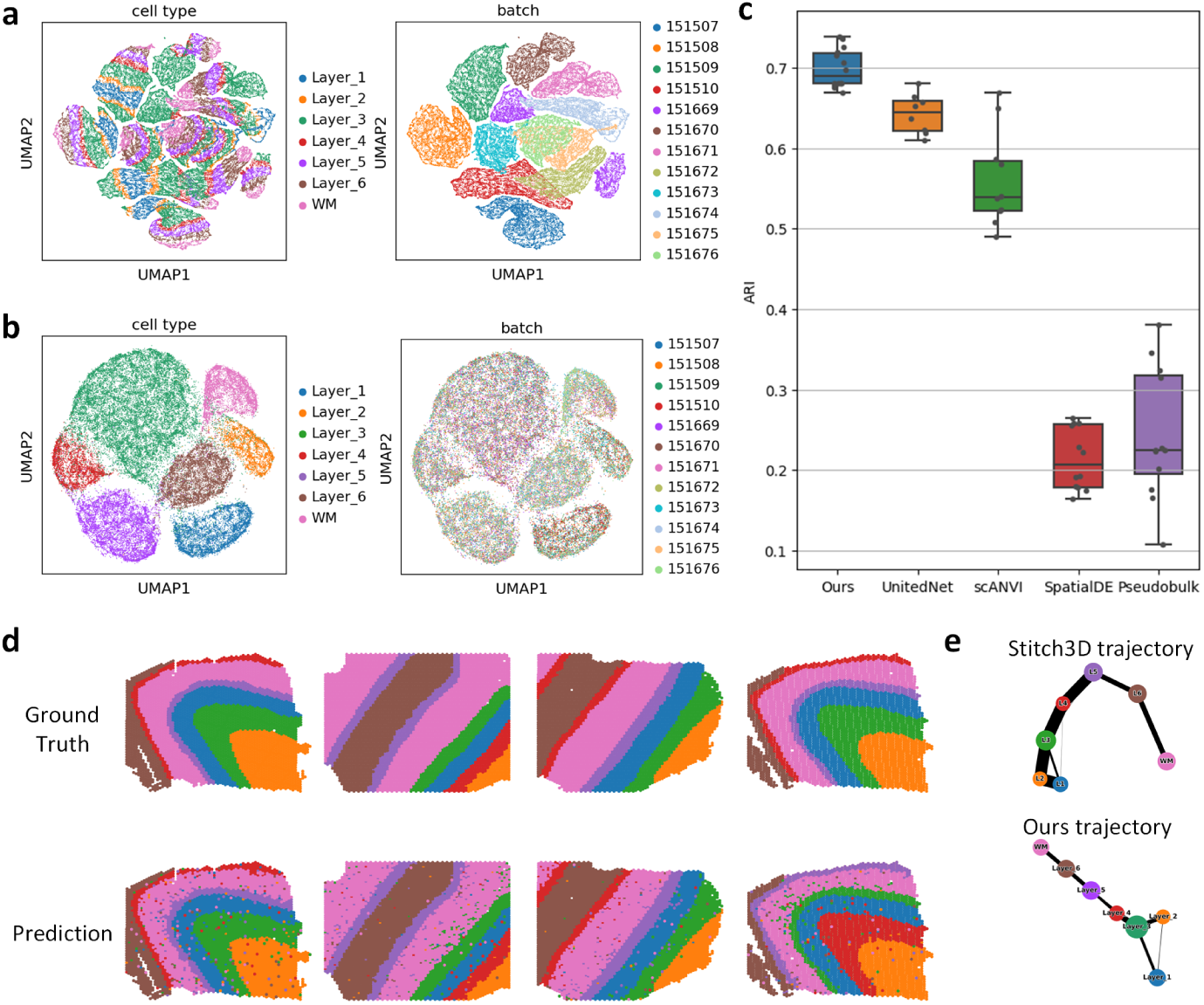
Applications on spatial data. **a**: UMAP visualization of multi-slice data after direct dimensionality reduction, colored by cell type and batch label respectively. **b**: UMAP visualization of cell embeddings output by scProto-Transformer, colored by cell type and batch label respectively. **c**: Comparison of results of different spatial data representation methods, we use ARI to measure the biological heterogeneity of cell embeddings. **d**: Visualization on slices, we compare the actual cell types with the predicted cell types. **e**: Comparison of trajectory inference results.

We further performed cell type reference mapping on the DLPFC dataset, which consists of 12 slices. We randomly selected 8 slices as the reference dataset and used the remaining 4 slices as the query dataset. As shown in Figure 5d, the predictions made by scProtoTransformer were highly consistent with the ground-truth annotations. Moreover, we compared our approach with Stitch3D [47], a more advanced 3D spatial transcriptomics method that leverages 3D spatial coordinates to accurately infer cell differentiation trajectories. Using scProtoTransformer’s cell embeddings combined with PAGA [48] for trajectory inference, we obtained consistent results (Figure 5e).

### 2.7 Donor level reference mapping

Multiple studies have demonstrated that multiple-instance learning (MIL) can effectively bridge donors and single-cell representations [49–51]. In MIL, a donor sample is treated as a bag, and the cells within the sample are treated as instances. Typically, the cancer phenotype of the donor sample (bag label) is known, while the phenotype of individual cells (instance labels) is unavailable. According to clinical diagnostic definitions, the relationship between bag and instance labels is as follows: if all instances in a bag are normal cells, the bag is defined as a normal bag; if a bag contains any tumor cells, it is defined as a tumor bag.

For a donor sample, we first obtain cell embeddings from scProtoTransformer and then input them into the MIL module, which aggregates them into a donor embedding. We then use donor-level embeddings for reference mapping, as shown in Figure 6a. By updating the MIL module based on the bag labels of reference donors, the module not only predicts the disease state of a donor but can also trace back—through the aggregation weights—which cells in the sample are malignant.

**Figure 6:**
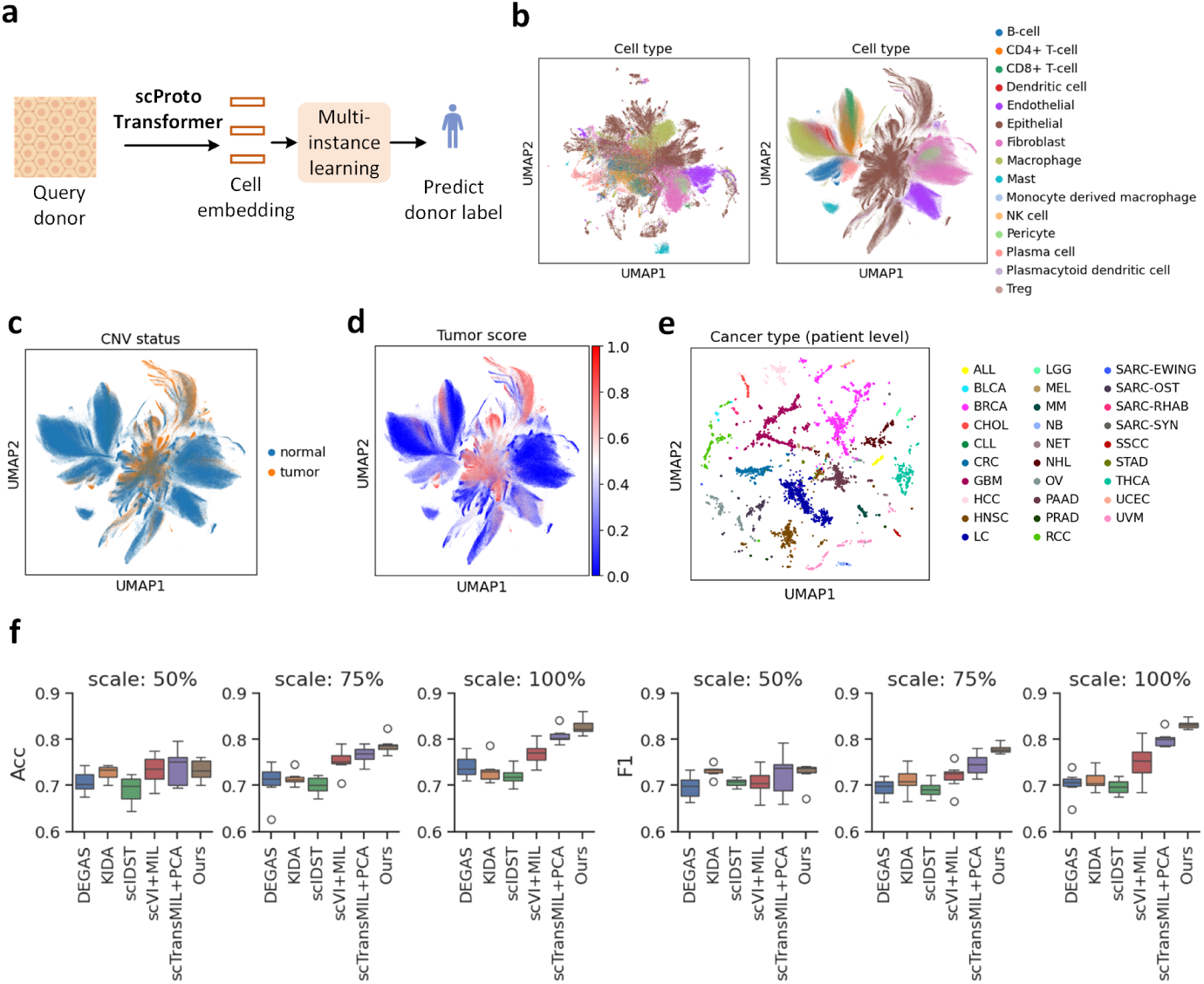
Donor level reference mapping. **a**: Workflow of donor-level reference mapping using scProtoTransformer. **b**: Visualization of cell embeddings on the Kang-2024 dataset, from left to right: PCA, scProtoTransformer, colored by cell type. **c**: Tumor labels at single-cell resolution. **d**: Tumor scores inferred by our method at single-cell resolution (our method can trace back from the donor level to the single-cell level). **e**: UMAP visualization at the donor level, colored by cancer type. **f**: Comparison of donor-level reference mapping results under different scales of training data.

We demonstrate this on the Kang-2024 dataset [52], which is composed of 104 publicly available cancer datasets. It covers 20,000 donor samples, 4 million single cells, and 30,000 genes. For reference mapping, we only use donor-level cancer labels (tumor or normal).

We visualized the PCA cell embeddings of Kang-2024 (colored by cell type) and also the cell embeddings generated by scProtoTransformer, as shown in Figure 6b. The comparison shows that clusters of different cell types are well separated, indicating that scProtoTransformer recovers the underlying biological heterogeneity.

In addition, we used the CNV labels provided in Kang-2024 as ground truth at single-cell resolution (Figure 6c). We then compared them with tumor scores at single-cell resolution inferred by the MIL module (Figure 6d). The inferred results are highly consistent with the ground truth. We also visualized donor-level embeddings with UMAP, colored by cancer type (Figure 6e), showing that our method effectively captures donor-level phenotypic heterogeneity.

For evaluation, we used Accuracy (Acc) and F1 score. The Kang-2024 dataset was randomly split into reference (80% of donors) and query (20% of donors) sets, ensuring no overlap of donors between the two, thereby directly assessing the model’s generalization to unseen donors. From the original reference set, we further sampled subsets of 50%, 75%, and 100% of donors. We trained on each subset and tested on the same query set. The performance of different methods is shown in Figure 6f. As the size of the reference data increased, all methods improved, but the improvement of scProtoTransformer was more significant, outperforming the baselines. This highlights the potential of scProtoTransformer for cancer detection at the donor level.

### 2.8 Ablation study

scProtoTransformer benefits from three key components: the knowledge-based prototype tokenizer, the knowledge distillation (KD) loss derived from the foundation model, and the dynamic SFT loss.

To systematically evaluate the contribution of each component, we conducted ablation experiments by individually removing the pathway-guided mask in the prototype tokenizer (w/o Proto), the KD loss (w/o Proto), and the dynamic SFT loss (w/o DFT), and then assessed performance changes across molecular, cell, and donor level tasks.

As shown in Table 1, removing the mask from the prototype tokenizer caused the most significant performance drop at the molecular level. This result confirms the critical role of the prototype tokenizer in mapping raw gene expression into biologically meaningful pathway space. It effectively filters out numerical batch noise while preserving and amplifying functional semantics of genes. In contrast, removing the KD loss or dynamic SFT loss had only a minor effect at the molecular level.

**Table 1:**
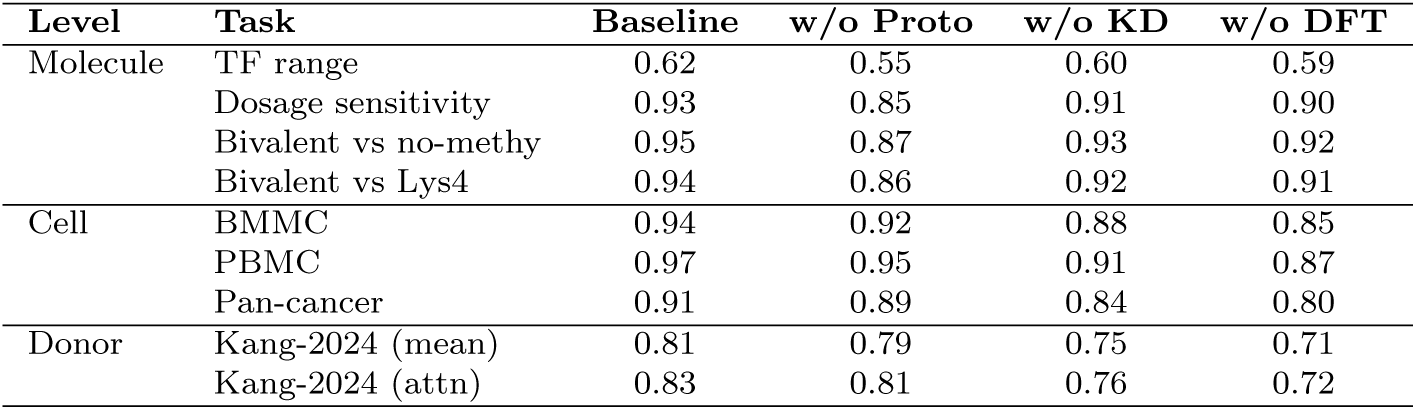
Ablation study results of scProtoTransformer across molecular, cell, and donor levels.

| Level | Task | Baseline | w/o Proto | w/o KD | w/o DFT |
| --- | --- | --- | --- | --- | --- |
| Molecule | TF range | 0.62 | 0.55 | 0.60 | 0.59 |
|  | Dosage sensitivity | 0.93 | 0.85 | 0.91 | 0.90 |
|  | Bivalent vs no-methy | 0.95 | 0.87 | 0.93 | 0.92 |
|  | Bivalent vs Lys4 | 0.94 | 0.86 | 0.92 | 0.91 |
| Cell | BMMC | 0.94 | 0.92 | 0.88 | 0.85 |
|  | PBMC | 0.97 | 0.95 | 0.91 | 0.87 |
|  | Pan-cancer | 0.91 | 0.89 | 0.84 | 0.80 |
| Donor | Kang-2024 (mean) | 0.81 | 0.79 | 0.75 | 0.71 |
|  | Kang-2024 (attn) | 0.83 | 0.81 | 0.76 | 0.72 |

At the cell and donor levels, however, removing the KD loss or dynamic SFT loss resulted in marked performance degradation. This indicates that the KD loss transfers the generalization ability of the foundation model through knowledge distillation, while the dynamic SFT loss enhances discriminability and robustness under limited labeled data by adaptively controlling gradients and preventing overfitting. This effect is further amplified in donor-level tasks. Overall, the ablation study validates the necessity of all three modules in scProtoTransformer: the prototype tokenizer drives molecular-level semantic modeling, while the KD loss and dynamic SFT loss jointly optimize discriminative representations at the cell and donor levels. Together, these components support cross-scale reference mapping.

## 3 Discussion

Single-cell omics analysis is entering the era of foundation models capable of integrating knowledge across diverse datasets. Despite significant progress made by large-scale foundation models, two major bottlenecks remain: persistent batch effects and the absence of a scalable, multi-level representation framework spanning molecular, cellular, and donor resolutions. We address these challenges with scProtoTransformer, a prototype-based Transformer that provides a unified, efficient, and interpretable solution for cross-scale reference mapping.

Instead of relying on raw expression values that are inherently noisy and sensitive to numerical variation, scProtoTransformer abstracts these signals into a set of stable functional prototypes defined by biological priors (pathways). This transformation effectively acts as a biological normalization layer, mitigating batch-induced “numerical drift” while preserving biologically meaningful patterns. The subsequent Transformer layers model interactions between these functional prototypes, producing cell embeddings that exhibit enhanced biological heterogeneity and reduced technical variation, as validated by our performance on multi-modal and batch integration tasks.

In addition, the stable training of scProtoTransformer benefits from our knowledge distillation strategy. By leveraging frozen large-scale foundation models (e.g., scGPT) as teachers, we distill high-quality representations into scProtoTransformer, while dynamic supervised fine-tuning (dynamic SFT) ensures biologically consistent gradient updates when labels are available. This allows scProtoTransformer to inherit the representational power of foundation models without requiring resource-intensive pretraining, making it easier for researchers to adapt foundation model knowledge into customized models.

scProtoTransformer systematically unifies representation learning across molecular, cell, and donor levels. Learned gene embeddings naturally cluster by function, enabling high-quality molecular-level reference mapping. At the cellular level, scProtoTransformer builds joint cell representations that facilitate integration and annotation, with prototype attention providing intrinsic interpretability by linking cell identities to specific active pathways. At the donor level, aggregating cell embeddings yields donor representations suitable for patient stratification and disease monitoring, while the aggregation weights allow inference of malignant or disease-relevant cell populations within a donor. Comprehensive benchmarks across multiple levels demonstrate that scProtoTransformer has the potential to serve as a unified framework for scalable reference mapping, thereby laying the foundation for systematic understanding of biological systems from genes to individuals.

## 4 Methods

### 4.1 Prototype tokenizer

The Prototype Tokenizer initializes model parameters with biological priors guided by known biological pathways. This initialization ensures that the learned prototypes correspond to biologically meaningful representations rather than purely numerical patterns.

The process begins by constructing a gene vocabulary (Gene Vocab) *W*_0_ ∈ ℝ*^n^*^×^*^h^*, where each row represents the embedding of a gene. The embedding *E_g_* ∈ ℝ*^h^* of each gene *g* is computed by summing the embeddings of the proteins encoded by the gene:

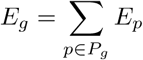

where *P_g_*is the set of proteins associated with gene *g*, and *E_p_* ∈ ℝ*^h^* is the embedding for protein *p* from ESM2 [53].

Next, a pathway-guided mask is applied to the Gene Vocab. This mask, based on known biological pathways, ensures that the model focuses attention on biologically relevant gene sets. The mask transforms the Gene Vocab into multiple parallel linear weights, each of which projects the gene expression into a prototype space.

For the knowledge-based mask, we create a list consisting of all gene names in reference dataset, and obtain *m* pathways corresponding to this gene list through GSEA (Gene Set Enrichment Analysis) combined with pathway annotation (gmt file on the web or locally). We added mask to Gene Vocab (a linear matrix) according to the relationship between genes and pathways. Assuming that gene 1 is not in pathway 1, the first row (gene 1) of linear matrix *W*_1_ ∈ ℝ*^n^*^×^*^h^* (pathway 1) is all masked to value 0. On the contrary, assuming that gene *n* is in pathway 1, the *n*-th row of linear matrix *W*_1_ ∈ ℝ*^n^*^×^*^h^* will not have a mask added. Finally, we get *m* matrices with masks (a parallel linear layer).

The parallel linear layer projects gene expression data into *m* prototypes, with each prototype token representing a specific biological pathway or function. The transformation from gene expression *x_i_* for a given cell *i* to prototype tokens *v_p_*is performed by applying the matrix of prototype *p*:

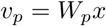

where *W_p_* is a masked matrix. After passing through *m* prototypes simultaneously, the result is a set of prototype tokens (*v*_1_*, …, v_m_*) representing biologically relevant functions.

### 4.2 scProtoTransformer architecture

The scProtoTransformer architecture models the interactions between the learned prototype tokens using 6 transformer layers with Flash Attention [54]. Once the prototype tokens are generated, a special `[CLS]` token is prepended to the sequence of prototype tokens. This token will eventually represent the embedding of the entire cell. The sequence, consisting of the `[CLS]` token followed by the prototype tokens, is then input into 6 stacked transformer layers.

The transformer layers in scProtoTransformer use Flash Attention, an optimized version of the attention mechanism that reduces memory usage and increases computation speed. Flash Attention performs the self-attention operation with reduced memory overhead by using memory-efficient techniques such as out-of-place operations. This enables the model to scale efficiently, even with large input sequences.

After passing through the 6 transformer layers, the final output is a sequence of embeddings, with the `[CLS]` token containing the most informative representation of the cell. This `[CLS]` token embedding is used as the cell embedding, which represents the biological state of the entire cell based on the learned prototype tokens and their interactions.

### 4.3 Foundation model distillation

We designed a knowledge distillation strategy guided by a foundation model for scProtoTransformer. The purpose of establishing this strategy is to enable scProtoTransformer to acquire the knowledge that the foundation model gains at a huge training cost, without the need for large-scale pre-training. Specifically, we use a frozen foundation model (such as scGPT) as a navigator, and supervise the learning of scProtoTransformer through the cell embedding of the navigator. Different from the unsupervised pre-training of previous foundation models, we can dynamically utilize the labels in the data. When there are available labels in the data, we leverage the discriminative of the labels themselves to enhance the biological representation capability of scProtoTransformer. To train the model on data of limited scale, we use dynamic SFT (Supervised Fine-Tuning) [55], which can adaptively control gradients to prevent the model from overfitting. The training objective consists of KD (knowledge distillation) loss and dynamic SFT. We first use KD loss on unlabeled data, and then use dynamic SFT on labeled data.

#### KD Loss

We use the foundation model scGPT as the teacher and scProtoTransformer as the student. We freeze the teacher and update the direction of the student’s output embedding based on cosine similarity. The cell embedding output by the teacher model is *c_t_* ∈ ℝ*^h^*, and the cell embedding output by the student model is *c_r_* ∈ ℝ*^h^*. First, we perform L2 normalization on both embeddings to eliminate the vector modulus and retain only the direction:

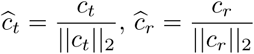

Then, we define the KD loss using cosine similarity:

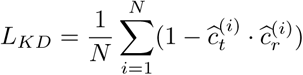

Where *N* is the total number of cells in the batch. By maximizing the similarity between the teacher and student embeddings of the same cell, the student model is able to quickly align the knowledge acquired by the foundation model during the large-scale pre-training phase.

#### Dynamic SFT

After scProtoTransformer is aligned using the KD loss, we use dynamic SFT to enhance the model’s discriminability. Standard SFT uses the cross-entropy loss. Dynamic SFT is a reinforcement learning-based SFT that avoids exploding gradients and prevents catastrophic forgetting caused by SFT. Let the model be the policy *π_θ_*, *θ* be the model parameters, the input sample be *x*, the ground truth label be *y*^∗^, and the model’s predicted label be *y*. For distribution *D*, the expectation of the SFT is:

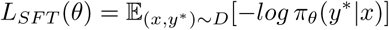

The gradient of *θ* is:

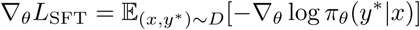

Through importance sampling, the SFT gradient is written as the policy gradient form used in reinforcement learning:

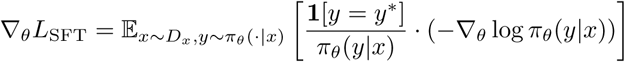

The above form exposes a key issue with SFT: when the model’s initial probability of the correct answer *π_θ_*(*y*^∗^|*x*) is low, the importance weight 1*/π_θ_*(*y*^∗^|*x*) can become very large, leading to exploding gradients and unstable training. To eliminate the unstable weight 1*/π_θ_*(*y*^∗^|*x*), we directly multiply the raw SFT gradient by a stop-grounded probability *sg*(*π_θ_*(*y*^∗^|*x*)) to offset it. Therefore, the dynamic SFT is:

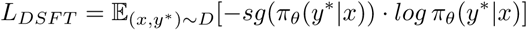

For a detailed derivation of the policy gradient, please refer to the supplementary material.

### 4.4 Scalable reference mapping

The scProtoTransformer framework supports scalable reference mapping at three biological levels: molecular, cellular, and donor levels.

#### Molecular level

The gene embeddings learned by scProtoTransformer can not only effectively capture the latent biological information of genes but also accurately assign labels to genes through supervised classification methods. This approach offers high flexibility, allowing extension to different cell types or biological conditions, and is suitable for various gene function prediction tasks.

In molecular level reference mapping, we leverage the gene vocabulary within scProtoTransformer to extract the embedding representation of each gene. Specifically, the gene vocabulary contains embedding vectors of multiple genes, which are learned through the scProtoTransformer model. To perform effective reference mapping on these gene embeddings, we adopt two common supervised learning methods: Logistic Regression (LR) and Random Forest (RF). These methods can learn the mapping relationship between genes and known labels in the gene embedding space. First, we construct an embedding representation for each gene. Then, we use known gene label data (such as the functional category of a gene or its expression status under certain biological conditions) to train a classifier. After training, the classifier can perform inference on unlabeled query genes and output predicted labels. These predicted labels help us accurately map the query genes to corresponding biological categories or functional groups, thereby achieving effective annotation of gene functions.

#### Cell level

When a single cell is input, scProtoTransformer can output a cell embedding. A cell classifier is used to assign predicted labels to each cell. We ensure that cell type annotations are consistent with the cell ontology [56, 57], enabling the classifier to predict the cell type for any input cell. If we want the model to be more adaptable to a specific dataset, we can fine-tune scProtoTransformer on that particular dataset.

scProtoTransformer accepts gene expression data as input. When dealing with scATAC-seq data, we use Seurat [31] to convert the peaks of scATAC-seq into genes corresponding to scRNA-seq. For single-modal data, scProtoTransformer has a built-in batch integration function, we only need to infer cell embeddings to obtain a cross-batch joint space. For multi-modal data, we first infer cell embeddings from the single-modal data, and then use an embedding alignment strategy [58] to integrate the cell embeddings across different modalities.

#### Donor level

In the donor level reference mapping, we use a multiple instance learning (MIL) module [59] to aggregate cell embeddings into a single donor level embedding. In multiple instance learning, a set of cells from a single donor forms a bag, while each individual cell is treated as an instance. Typically, the bag label is known (the donor label), while the instance labels are unavailable. The goal is to learn a model that aggregates cell embeddings within each bag to form a donor level embedding and uses this embedding for donor label prediction.

For this task, cell embeddings are derived from the scProtoTransformer model, which represents each cell in a high-dimensional feature space. Given a donor with *N* cells, the embedding of the *i*-th cell is denoted as **e***_i_*, where *i* = 1, 2, …, *N*. The objective is to learn an aggregation function that combines these cell embeddings into a single donor embedding **e***_donor_* and to predict the donor’s label based on this embedding.

We adopt an attention-based multiple instance learning approach. Specifically, we assign an attention score *α_i_* to each cell embedding **e***_i_*, which reflects how relevant that cell is for the donor label. The attention scores are computed via a dense layer, where higher scores indicate a stronger correlation with the donor label.

Next, we aggregate the cell embeddings to form a donor level embedding **e***_donor_*. The aggregation is performed as a weighted sum of the individual cell embeddings, where the weight for each cell embedding is its corresponding attention score *α_i_*. The aggregation function is defined as:

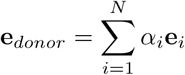

This weighted sum ensures that cells with higher attention scores contribute more to the final donor level embedding **e***_donor_*. Once we have the aggregated donor embedding **e***_donor_*, we can use a classifier to predict the donor’s label. In addition to label prediction, we can also backtrack to the instances that contribute most to the donor label prediction. Since attention scores *α_i_* indicate the relevance of each cell to the donor label, higher attention scores correspond to cells that are more important for the prediction.

### 4.5 Experimental setting

In scProtoTransformer, we followed TOSICA and KIDA, the knowledge database required by scProtoTransformer when generating functional patterns comes from [60, 61], and we set *m*=300. Enrichment analysis tool is GSEApy [62], preprocessing and differential analysis tool is scanpy [63]. scProtoTransformer is trained using NVIDIA GeForce RTX A6000 with 48 GB memory. Adam [64] optimizer with 1e-4 learning rate is used to update model parameters. The batch size is set to 16.

We sampled 2 million cells from the human subset of scBaseCamp [23] and searched annotation information for each cell using scsimilarity [57], filtering out low-confidence labels. scProtoTransformer first aligned the cell embeddings output by scGPT across all cells. We then performed dynamic SFT on the model using the labeled data.

## Data availability

All raw datasets used in this study are already published and were obtained from public data repositories.

## Code availability

The code of this study is available at https://github.com/shapsider/ scprototransformer.

## Acknowledgments

This work was supported by the National Key Research and Development Program of China (No. 2024YFA0917600, to C.Y.-C.C.).

## Author Contributions

Z.T. and C.Y.-C.C. designed research. Z.T., H.H., G.C., S.C., T.L., J.Z. and J.H. worked together to complete the experiment and analyze the data. L.Y. and C.Y.-C.C. contributed to analytic tools. Z.T., H.H. and G.C. wrote the manuscript.

## 5 Supplementary texts

### 5.1 Derivation process of dynamic SFT

The standard supervised fine-tuning (SFT) objective minimizes the negative log-likelihood of expert demonstrations:

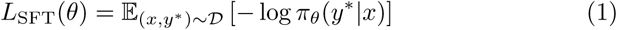

where *x* is input data. *y*^∗^ is expert label. *π_θ_*(*y*^∗^|*x*) is model’s likelihood of expert label given parameters *θ*. The gradient of this objective is:

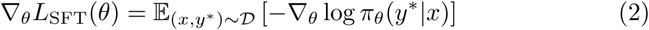

The reinforcement learning objective maximizes expected reward:

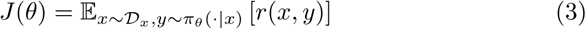

with the policy gradient theorem yielding:

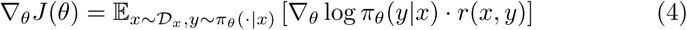

Applying importance sampling to the SFT gradient:

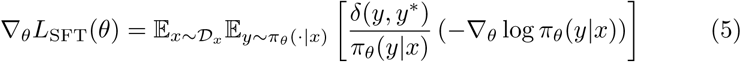

where *δ*(*y, y*^∗^) is the Kronecker delta function (1 when *y* = *y*^∗^, 0 otherwise). Defining implicit reward and importance weight functions:

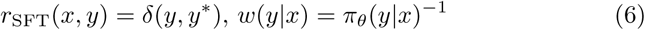

yields the RL-equivalent form:

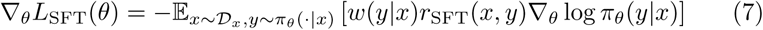

The importance weight *w*(*y*|*x*) = *π_θ_*(*y*^∗^|*x*)^−1^ introduces instability:

- Gradient explosion when *π_θ_*(*y*^∗^|*x*) → 0^+^.
- High-variance updates during early training phases.
- Degraded generalization due to overemphasis on low-likelihood labels.

Dynamic SFT (DFT) introduces a stabilizing coefficient *π_θ_*(*y*^∗^|*x*):

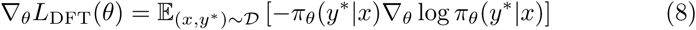

where *π_θ_*(*y*^∗^|*x*) is treated as a fixed scaling factor (stop-gradient).

The corresponding DFT loss function:

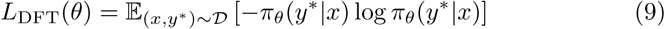

## 6 Supplementary figures

**Figure supplementary 1:**
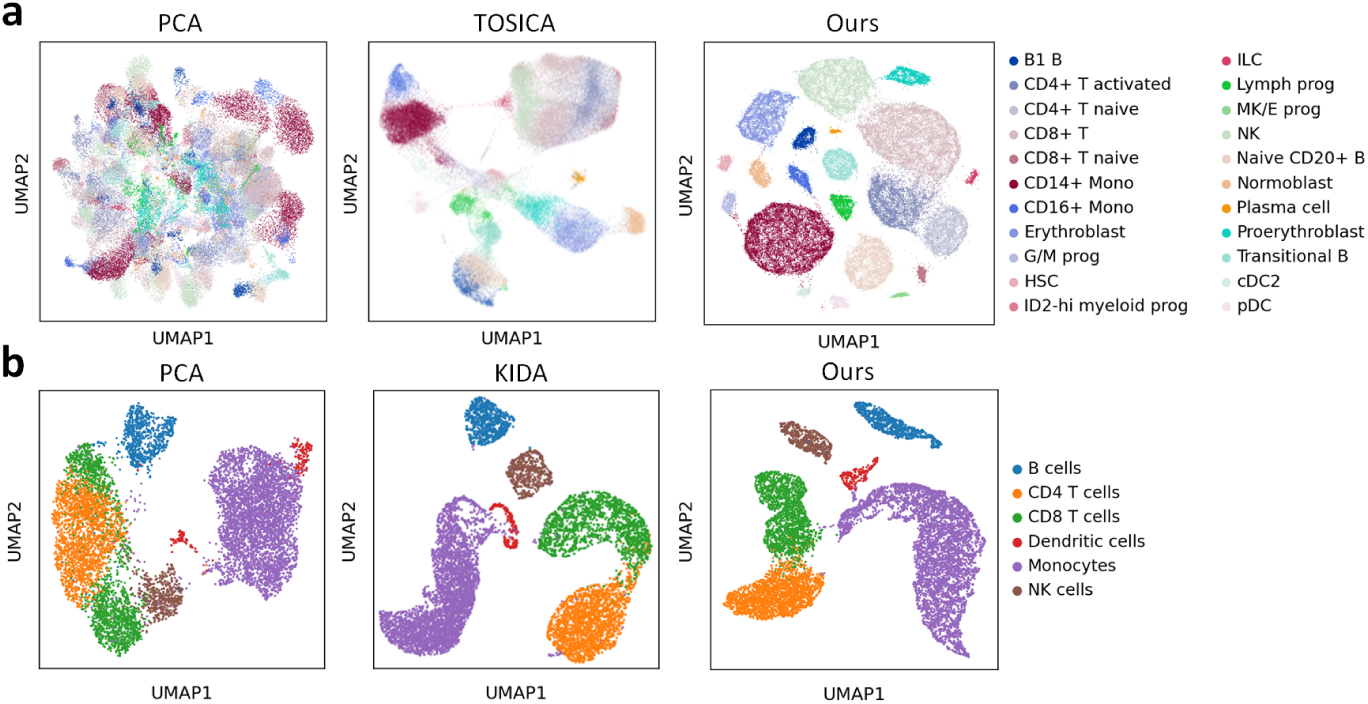
Cell-scale UMAP visualization. Figure a: Cell embedding UMAP of the BMMC dataset (measured using scRNA-seq). From left to right, PCA, TOSICA (a method specific to scRNA-seq), and our cell embeddings are visualized, colored by cell type. Figure b: Cell embedding UMAP of the PBMC dataset (measured using scATAC-seq). From left to right, PCA, KIDA (a method specific to scATAC-seq), and our cell embeddings are visualized, colored by cell type.

**Figure supplementary 2:**
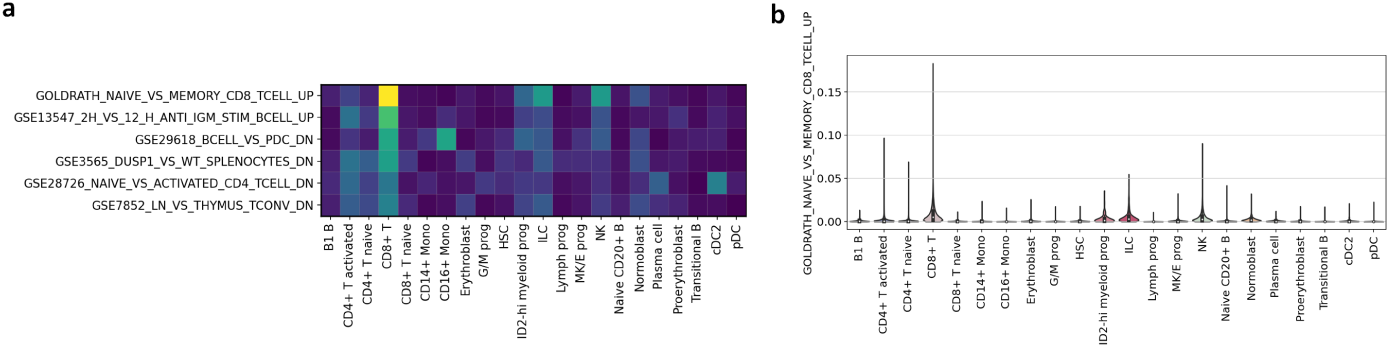
Results of interpretable attention on prototypes. Figure a: The vertical axis represents the top six prototypes specific to CD8+ T cells, and the horizontal axis represents different cell types, with attention visualized using a heatmap. Figure b: For a given prototype (the first prototype in Figure a, named by pathway), the gene expression distribution corresponding to the prototype is calculated across different cell types.

**Figure supplementary 3:**
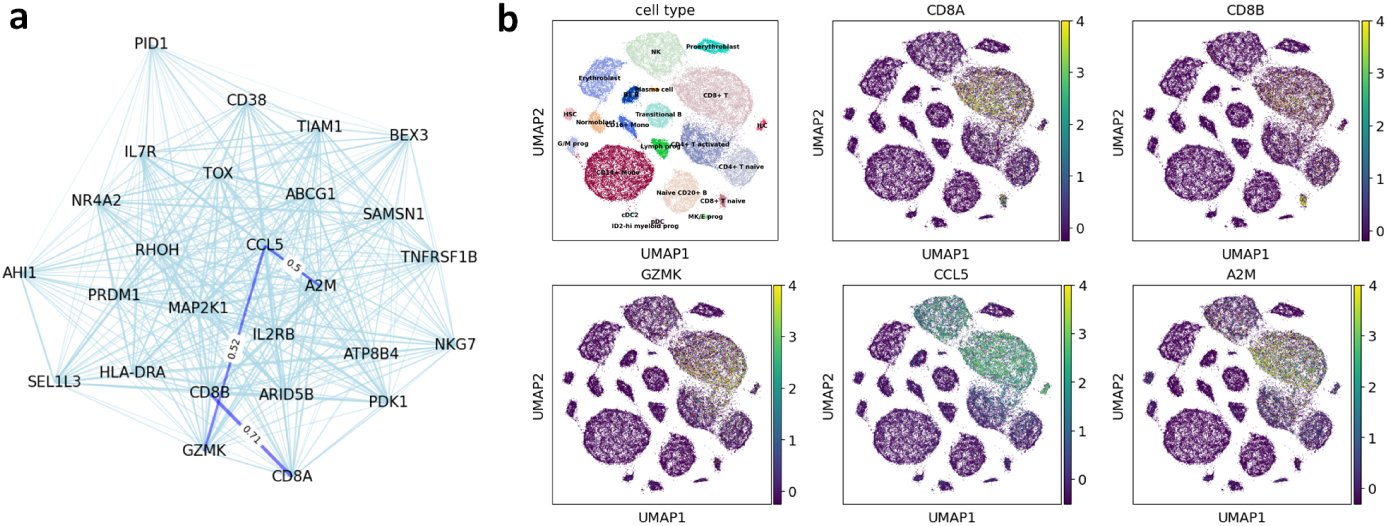
Gene interpretability. Figure a: The CD8+ T-specific gene co-expression network from scProtoTransformer. The edges with high similarity values are shown in dark blue. Figure b: Validation results of important genes based on UMAP visualization.

**Figure supplementary 4:**
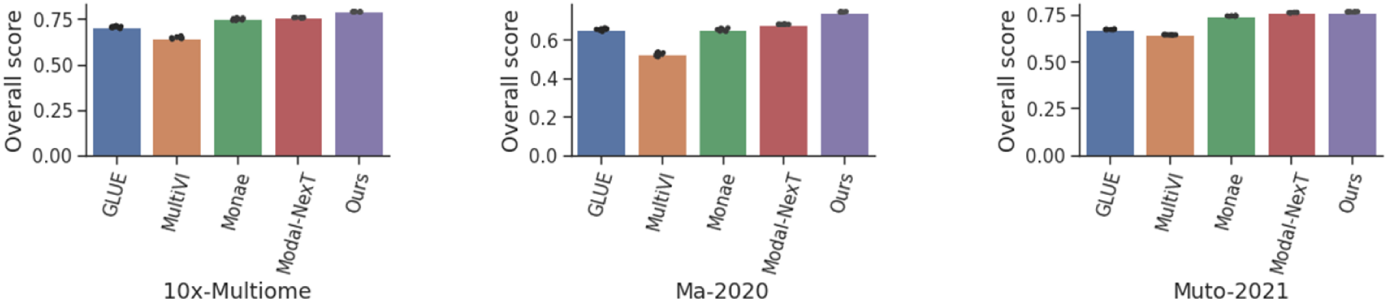
Overall score comparison of multi-modal integration.

## References

[1] Rood, J.E., Wynne, S., Robson, L., Hupalowska, A., Randell, J., Teichmann, S.A., Regev, A.: The human cell atlas from a cell census to a unified foundation model. Nature 637(8048), 1065–1071 (2025)

[2] Lotfollahi, M., Hao, Y., Theis, F.J., Satija, R.: The future of rapid and automated single-cell data analysis using reference mapping. Cell 187(10), 2343–2358 (2024)

[3] He, H., Chen, G., Tang, Z., Chen, C.Y.-C.: Dual modality feature fused neural network integrating binding site information for drug target affinity prediction. NPJ Digital Medicine 8(1), 67 (2025)

[4] He, H., Tang, Z., Chen, G., Xu, F., Hu, Y., Feng, Y., Wu, J., Huang, Y.-A., Huang, Z.-A., Tan, K.C.: sckan: Interpretable single-cell analysis for cell-type-specific gene discovery and drug repurposing via kolmogorov-arnold networks. bioRxiv, 2025-02 (2025)

[5] Chen, S., Tang, Z., You, L., Chen, C.Y.-C.: A knowledge distillation-guided equivariant graph neural network for improving protein interaction site prediction performance. Knowledge-Based Systems 300, 112209 (2024)

[6] Tang, Z., Chen, G., Yang, H., Zhong, W., Chen, C.Y.-C.: Dsil-ddi: A domain-invariant substructure interaction learning for generalizable drug–drug interaction prediction. IEEE Transactions on Neural Networks and Learning Systems 35(8), 10552–10560 (2023)

[7] Bunne, C., Roohani, Y., Rosen, Y., Gupta, A., Zhang, X., Roed, M., Alexandrov, T., AlQuraishi, M., Brennan, P., Burkhardt, D.B., et al.: How to build the virtual cell with artificial intelligence: Priorities and opportunities. Cell 187(25), 7045–7063 (2024)

[8] Tang, Z., Huang, J., Chen, G., Chen, C.Y.-C.: Comprehensive view embedding learning for single-cell multimodal integration. In: Proceedings of the AAAI Conference on Artificial Intelligence, vol. 38, pp. 15292–15300 (2024)

[9] Zhu, J., Li, Y., Tang, Z., Chang, C.: Dusted: Dual-attention enhanced spatial transcriptomics denoiser. In: Proceedings of the AAAI Conference on Artificial Intelligence, vol. 39, pp. 1219–1227 (2025)

[10] Zhu, J., Zhang, Z., Xiang, Y., Xie, B., Dong, X., Xie, L., Zhou, P., Yao, R., Wang, X., Li, Y., et al.: Decoding cell identity with multi-scale explainable deep learning. bioRxiv, 2024–02 (2024)

[11] Tejada-Lapuerta, A., Bertin, P., Bauer, S., Aliee, H., Bengio, Y., Theis, F.J.: Causal machine learning for single-cell genomics. Nature Genetics, 1–12 (2025)

[12] Wang, Y., Thistlethwaite, W., Tadych, A., Ruf-Zamojski, F., Bernard, D.J., Cappuccio, A., Zaslavsky, E., Chen, X., Sealfon, S.C., Troyanskaya, O.G.: Automated single-cell omics end-to-end framework with data-driven batch inference. Cell Systems 15(10), 982–990 (2024)

[13] Theodoris, C.V., Xiao, L., Chopra, A., Chaffin, M.D., Al Sayed, Z.R., Hill, M.C., Mantineo, H., Brydon, E.M., Zeng, Z., Liu, X.S., et al.: Transfer learning enables predictions in network biology. Nature 618(7965), 616–624 (2023)

[14] Yang, F., Wang, W., Wang, F., Fang, Y., Tang, D., Huang, J., Lu, H., Yao, J.: scbert as a large-scale pretrained deep language model for cell type annotation of single-cell rna-seq data. Nature Machine Intelligence 4(10), 852–866 (2022)

[15] Hao, M., Gong, J., Zeng, X., Liu, C., Guo, Y., Cheng, X., Wang, T., Ma, J., Zhang, X., Song, L.: Large-scale foundation model on single-cell transcriptomics. Nature Methods 21(8), 1481–1491 (2024)

[16] Cui, H., Wang, C., Maan, H., Pang, K., Luo, F., Duan, N., Wang, B.: scgpt: toward building a foundation model for single-cell multi-omics using generative ai. Nature Methods 21(8), 1470–1480 (2024)

[17] Zeng, Y., Xie, J., Shangguan, N., Wei, Z., Li, W., Su, Y., Yang, S., Zhang, C., Zhang, J., Fang, N., et al.: Cellfm: a large-scale foundation model pre-trained on transcriptomics of 100 million human cells. Nature Communications 16(1), 4679 (2025)

[18] Chen, H., Ryu, J., Vinyard, M.E., Lerer, A., Pinello, L.: Simba: single-cell embedding along with features. Nature Methods 21(6), 1003–1013 (2024)

[19] Cui, H., Tejada-Lapuerta, A., Brbić, M., Saez-Rodriguez, J., Cristea, S., Goodarzi, H., Lotfollahi, M., Theis, F.J., Wang, B.: Towards multimodal foundation models in molecular cell biology. Nature 640(8059), 623–633 (2025)

[20] Tang, Z., Yang, H., Chen, C.Y.-C.: Weakly supervised posture mining for fine-grained classification. In: Proceedings of the IEEE/CVF Conference on Computer Vision and Pattern Recognition, pp. 23735–23744 (2023)

[21] Li, Y., Tan, G., Gou, C.: Cascaded iterative transformer for jointly predicting facial landmark, occlusion probability and head pose. International Journal of Computer Vision 132(4), 1242–1257 (2024)

[22] Li, Y., Wang, S., Tan, G.: Id-nerf: Indirect diffusion-guided neural radiance fields for generalizable view synthesis. Expert Systems with Applications 266, 126068 (2025)

[23] Youngblut, N.D., Carpenter, C., Prashar, J., Ricci-Tam, C., Ilango, R., Teyssier, N., Konermann, S., Hsu, P.D., Dobin, A., Burke, D.P., et al.: scbasecount: an ai agent-curated, uniformly processed, and continually expanding single cell data repository. bioRxiv, 2025-02 (2025)

[24] McInnes, L., Healy, J., Melville, J.: Umap: Uniform manifold approximation and projection for dimension reduction. arXiv preprint arXiv:1802.03426 (2018)

[25] Chen, Y., Zou, J.: Simple and effective embedding model for single-cell biology built from chatgpt. Nature biomedical engineering 9(4), 483–493 (2025)

[26] Luecken, M.D., Burkhardt, D.B., Cannoodt, R., Lance, C., Agrawal, A., Aliee, H., Chen, A.T., Deconinck, L., Detweiler, A.M., Granados, A.A., et al.: A sandbox for prediction and integration of dna, rna, and proteins in single cells. In: Thirty-fifth Conference on Neural Information Processing Systems Datasets and Benchmarks Track (Round 2) (2021)

[27] Satpathy, A.T., Granja, J.M., Yost, K.E., Qi, Y., Meschi, F., McDermott, G.P., Olsen, B.N., Mumbach, M.R., Pierce, S.E., Corces, M.R., et al.: Massively parallel single-cell chromatin landscapes of human immune cell development and intratumoral t cell exhaustion. Nature biotechnology 37(8), 925–936 (2019)

[28] Liu, T., Wang, Y., Ying, R., Zhao, H.: Muse-gnn: Learning unified gene representation from multimodal biological graph data. Advances in neural information processing systems 36, 24661–24677 (2023)

[29] Zheng, L., Qin, S., Si, W., Wang, A., Xing, B., Gao, R., Ren, X., Wang, L., Wu, X., Zhang, J., et al.: Pan-cancer single-cell landscape of tumorinfiltrating t cells. Science 374(6574), 6474 (2021)

[30] Cheng, S., Li, Z., Gao, R., Xing, B., Gao, Y., Yang, Y., Qin, S., Zhang, L., Ouyang, H., Du, P., et al.: A pan-cancer single-cell transcriptional atlas of tumor infiltrating myeloid cells. Cell 184(3), 792–809 (2021)

[31] Stuart, T., Butler, A., Hoffman, P., Hafemeister, C., Papalexi, E., Mauck, W.M., Hao, Y., Stoeckius, M., Smibert, P., Satija, R.: Comprehensive integration of single-cell data. cell 177(7), 1888–1902 (2019)

[32] Domínguez Conde, C., Xu, C., Jarvis, L., Rainbow, D., Wells, S., Gomes, T., Howlett, S., Suchanek, O., Polanski, K., King, H., et al.: Cross-tissue immune cell analysis reveals tissue-specific features in humans. Science 376(6594), 5197 (2022)

[33] Ma, F., Pellegrini, M.: Actinn: automated identification of cell types in single cell rna sequencing. Bioinformatics 36(2), 533–538 (2020)

[34] Chen, J., Xu, H., Tao, W., Chen, Z., Zhao, Y., Han, J.-D.J.: Transformer for one stop interpretable cell type annotation. Nature Communications 14(1), 223 (2023)

[35] Ma, W., Lu, J., Wu, H.: Cellcano: supervised cell type identification for single cell atac-seq data. Nature Communications 14(1), 1864 (2023)

[36] Zhang, Y., Xiang, G., Jiang, A.Y., Lynch, A., Zeng, Z., Wang, C., Zhang, W., Fan, J., Kang, J., Gu, S.S., et al.: Metatime integrates single-cell gene expression to characterize the meta-components of the tumor immune microenvironment. Nature communications 14(1), 2634 (2023)

[37] Zhao, S., Zhang, J., Nie, Z.: Large-scale cell representation learning via divide-and-conquer contrastive learning. arXiv preprint arXiv:2306.04371 (2023)

[38] Zhao, S., Zhang, J., Luo, Y., Wu, Y., Nie, Z.: Langcell: Language-cell pre-training for cell identity understanding. arXiv preprint arXiv:2405.06708 (2024)

[39] Tang, Z., Chen, G., Chen, S., He, H., You, L., Chen, C.Y.-C.: Knowledge-based inductive bias and domain adaptation for cell type annotation. Communications biology 7(1), 1440 (2024)

[40] Zhang, X., Lan, Y., Xu, J., Quan, F., Zhao, E., Deng, C., Luo, T., Xu, L., Liao, G., Yan, M., et al.: Cellmarker: a manually curated resource of cell markers in human and mouse. Nucleic acids research 47(D1), 721–728 (2019)

[41] Tang, Z., Chen, G., Chen, S., Yao, J., You, L., Chen, C.Y.-C.: Modal-nexus auto-encoder for multi-modality cellular data integration and imputation. Nature Communications 15(1), 9021 (2024)

[42] Cao, Z.-J., Gao, G.: Multi-omics single-cell data integration and regulatory inference with graph-linked embedding. Nature Biotechnology 40(10), 1458–1466 (2022)

[43] Ma, S., Zhang, B., LaFave, L.M., Earl, A.S., Chiang, Z., Hu, Y., Ding, J., Brack, A., Kartha, V.K., Tay, T., et al.: Chromatin potential identified by shared single-cell profiling of rna and chromatin. Cell 183(4), 1103–1116 (2020)

[44] Muto, Y., Wilson, P.C., Ledru, N., Wu, H., Dimke, H., Waikar, S.S., Humphreys, B.D.: Single cell transcriptional and chromatin accessibility profiling redefine cellular heterogeneity in the adult human kidney. Nature Communications 12(1), 2190 (2021)

[45] Ashuach, T., Gabitto, M.I., Koodli, R.V., Saldi, G.-A., Jordan, M.I., Yosef, N.: Multivi: deep generative model for the integration of multi-modal data. Nature Methods 20(8), 1222–1231 (2023)

[46] Tang, Z., Chen, G., Chen, S., He, H., Huang, J., Dong, T., Zhou, J., Zhao, L., You, L., Chen, C.Y.-C.: Modal-next: Toward unified heterogeneous cellular data integration. Information Fusion, 103479 (2025)

[47] Wang, G., Zhao, J., Yan, Y., Wang, Y., Wu, A.R., Yang, C.: Construction of a 3d whole organism spatial atlas by joint modelling of multiple slices with deep neural networks. Nature Machine Intelligence 5(11), 1200–1213 (2023)

[48] Wolf, F.A., Hamey, F.K., Plass, M., Solana, J., Dahlin, J.S., Göttgens, B., Rajewsky, N., Simon, L., Theis, F.J.: Paga: graph abstraction reconciles clustering with trajectory inference through a topology preserving map of single cells. Genome biology 20(1), 59 (2019)

[49] Tang, Z., Wang, F., Yang, F., Song, J., Chen, C.Y.-C., Yao, J.: sctransmil bridges patient-level phenotypes and single-cell transcriptomics for cancer screening and heterogeneity inference. bioRxiv, 2025–04 (2025)

[50] Litinetskaya, A., Shulman, M., Hediyeh-zadeh, S., Moinfar, A.A., Curion, F., Szalata, A., Omidi, A., Lotfollahi, M., Theis, F.J.: Multimodal weakly supervised learning to identify disease-specific changes in single-cell atlases. bioRxiv, 2024–07 (2024)

[51] Wehbe, F., Adams, L., Babadoudou, J., Yuen, S., Kim, Y.-S., Tanaka, Y.: Inferring disease progression stages in single-cell transcriptomics using a weakly supervised deep learning approach. Genome research 35(1), 135–146 (2025)

[52] Kang, J., Lee, J.H., Cha, H., An, J., Kwon, J., Lee, S., Kim, S., Baykan, M.Y., Kim, S.Y., An, D., et al.: Systematic dissection of tumor-normal single-cell ecosystems across a thousand tumors of 30 cancer types. Nature communications 15(1), 4067 (2024)

[53] Lin, Z., Akin, H., Rao, R., Hie, B., Zhu, Z., Lu, W., Smetanin, N., Verkuil, R., Kabeli, O., Shmueli, Y., et al.: Evolutionary-scale prediction of atomic-level protein structure with a language model. Science 379(6637), 1123–1130 (2023)

[54] Dao, T., Fu, D., Ermon, S., Rudra, A., Ré, C.: Flashattention: Fast and memory-efficient exact attention with io-awareness. Advances in neural information processing systems 35, 16344–16359 (2022)

[55] Wu, Y., Zhou, Y., Ziheng, Z., Peng, Y., Ye, X., Hu, X., Zhu, W., Qi, L., Yang, M.-H., Yang, X.: On the generalization of sft: A reinforcement learning perspective with reward rectification. arXiv preprint arXiv:2508.05629 (2025)

[56] Ergen, C., Xing, G., Xu, C., Kim, M., Jayasuriya, M., McGeever, E., Oliveira Pisco, A., Streets, A., Yosef, N.: Consensus prediction of cell type labels in single-cell data with popv. Nature Genetics 56(12), 2731–2738 (2024)

[57] Heimberg, G., Kuo, T., DePianto, D.J., Salem, O., Heigl, T., Diamant, N., Scalia, G., Biancalani, T., Turley, S.J., Rock, J.R., et al.: A cell atlas foundation model for scalable search of similar human cells. Nature 638(8052), 1085–1094 (2025)

[58] Korsunsky, I., Millard, N., Fan, J., Slowikowski, K., Zhang, F., Wei, K., Baglaenko, Y., Brenner, M., Loh, P.-r., Raychaudhuri, S.: Fast, sensitive and accurate integration of single-cell data with harmony. Nature Methods 16(12), 1289–1296 (2019)

[59] Ilse, M., Tomczak, J., Welling, M.: Attention-based deep multiple instance learning. In: International Conference on Machine Learning, pp. 2127–2136 (2018). PMLR

[60] Subramanian, A., Tamayo, P., Mootha, V.K., Mukherjee, S., Ebert, B.L., Gillette, M.A., Paulovich, A., Pomeroy, S.L., Golub, T.R., Lander, E.S., et al.: Gene set enrichment analysis: a knowledge-based approach for interpreting genome-wide expression profiles. Proceedings of the National Academy of Sciences 102(43), 15545–15550 (2005)

[61] Mootha, V.K., Lindgren, C.M., Eriksson, K.-F., Subramanian, A., Sihag, S., Lehar, J., Puigserver, P., Carlsson, E., Ridderstråle, M., Laurila, E., et al.: Pgc-1*α*-responsive genes involved in oxidative phosphorylation are coordinately downregulated in human diabetes. Nature genetics 34(3), 267–273 (2003)

[62] Fang, Z., Liu, X., Peltz, G.: Gseapy: a comprehensive package for performing gene set enrichment analysis in python. Bioinformatics 39(1), 757 (2023)

[63] Wolf, F.A., Angerer, P., Theis, F.J.: Scanpy: large-scale single-cell gene expression data analysis. Genome biology 19(1), 15 (2018)

[64] Kingma, D.P.: Adam: A method for stochastic optimization. arXiv preprint arXiv:1412.6980 (2014)

